# Benchmarking algorithms for genomic prediction of complex traits

**DOI:** 10.1101/614479

**Authors:** Christina B. Azodi, Andrew McCarren, Mark Roantree, Gustavo de los Campos, Shin-Han Shiu

## Abstract

The usefulness of Genomic Prediction (GP) in crop and livestock breeding programs has led to efforts to develop new and improved GP approaches including non-linear algorithm, such as artificial neural networks (ANN) (i.e. deep learning) and gradient tree boosting. However, the performance of these algorithms has not been compared in a systematic manner using a wide range of GP datasets and models. Using data of 18 traits across six plant species with different marker densities and training population sizes, we compared the performance of six linear and five non-linear algorithms, including ANNs. First, we found that hyperparameter selection was critical for all non-linear algorithms and that feature selection prior to model training was necessary for ANNs when the markers greatly outnumbered the number of training lines. Across all species and trait combinations, no one algorithm performed best, however predictions based on a combination of results from multiple GP algorithms (i.e. ensemble predictions) performed consistently well. While linear and non-linear algorithms performed best for a similar number of traits, the performance of non-linear algorithms vary more between traits than that of linear algorithms. Although ANNs did not perform best for any trait, we identified strategies (i.e. feature selection, seeded starting weights) that boosted their performance near the level of other algorithms. These results, together with the fact that even small improvements in GP performance could accumulate into large genetic gains over the course of a breeding program, highlights the importance of algorithm selection for the prediction of trait values.

## Introduction

The ability to predict complex traits from genotypes is a grand challenge in biology and is accelerating the speed of crop and livestock breeding (1–4). Genomic Prediction (GP, aka Genomic Selection), the use of genome-wide genetic markers to predict complex traits, was originally proposed by Meuwissen *et al.* (5) as a solution to the limitations of Marker-Assisted Selection where only a limited number of previously identified markers with the strongest associations are used for breeding value prediction. GP is particularly well-suited for the prediction of quantitative traits controlled by many small-effect alleles (6). A major challenge in using GP is estimating the effects of a large number of makers (p) using phenotype information of a comparatively limited number of individuals (n) (i.e. p >> n) (5). To address this challenge, Meuwissen *et al.* (5) first presented three statistical methods for GP. The first is a linear mixed model called ridge regression Best Linear Unbiased Prediction (rrBLUP), which uniformly shrinks the marker effects. The other two are Bayesian approaches, BayesA (BA) and BayesB (BB), which both differentially shrink the marker effects and with BB also performing variable selection. Since then, additional approaches have been shown to be useful for GP, including Least Absolute Angle and Selection Operator (LASSO) (7), Elastic Net (8), Support Vector Regression with a linear kernel (SVR_lin_) (9,10), and additional Bayesian methods including Bayesian LASSO (BL), BayesCπ, and BayesDπ (11,12).

While these approaches perform well when dealing with high dimensional data (i.e. p>>n), they are all based on a linear mapping from genotype to phenotypes, and therefore may not fully capture non-linear effects (i.e. epistasis, dominance), which are likely to be important for complex traits (13,14). To overcome this limitation, non-linear approaches have been applied to GP problems, including reproducing kernel Hilbert spaces regression (15,16), Support Vector Regression with non-linear kernels (i.e. polynomial SVR_poly_ and radial basis function SVR_rbf_ (17,18), and decision tree based algorithms such as Random Forest (RF) (19,20) and Gradient Tree Boosting (GTB) (21). In previous efforts comparing the performance of multiple linear and non-linear approaches (22–26), no single method performs best in all cases. Factors such as the size of the training data set, marker type and number, trait heritability, effective population size, the number of causal loci, as well as genetic architecture (the locus effect size distribution) can all affect algorithm performance (20,27–29).This highlights the importance of comparing new algorithms across a diverse range of datasets and models.

With improvements in computing speeds, the development of graphics processing units (GPUs), and breakthroughs in algorithms for backpropagation learning (30,31), there has been a resurgence of research using artificial neural networks (ANNs) to model complex biological processes (32,33). ANNs are a class of machine learning methods that perform layers of transformations on features to create abstraction features, known as hidden layers, which are used for predictions. The first application of ANNs for GP was presented in 2011, when Okut *et al.* built fully connected ANNs (i.e. each node in a layer is connected to all nodes in surrounding layers, also called a multilayer perceptron) containing one hidden layer to predict body mass index in mice (34). Since 2011, more complex ANN architectures have been used for GP including radial basis function neural networks (35) and deep neural networks (36,37), deep recurrent neural networks (38), probabilistic neural network classifiers (39,40), and convolutional neural networks (41). As sequencing continues to become less expensive, whole-genome marker datasets are becoming larger, with some breeding programs generating data for hundreds of thousands of markers. Because of the internal complexity of ANN models, training an ANN with so many markers can result in sub-optimal solutions or underfitting. Therefore, it is especially important to benchmark ANNs against other GP statistical approaches on high dimensionality datasets.

While GP studies using a variety of algorithms have yielded promising results, most have focused on a small number of datasets. In addition, a comprehensive comparison of GP algorithms, particularly ANNs, on a wide range of GP problems is missing (**Figure 1A**). Here we compared the ability of 11 GP algorithms (see **Methods**, **Figure 1B**) to predict a diverse range of physiological traits in six plant species (maize, rice, sorghum, soy, spruce, and switchgrass; **Figure 1C**). These six data sets (referred to as the benchmark data sets) represent a wide range of GP data types, with the size of the training data set ranging from 327 to 5,014 individuals, and 4,000 to 332,000 markers derived from array-based approaches or sequencing. Compared to the linear algorithms included in the study, the non-linear algorithms, especially ANNs, require more pre-modeling tuning (e.g. hyperparameter selection, feature selection). Therefore, before comparing algorithm performance across all 18 combinations of species and traits, we first focused on predicting plant height in each species in order to establish best practices for model building. Because ANNs are underrepresented in GP comparison studies and our first attempts to use ANNs for GP performed relatively poorly, we focus on methods to improve ANN performance, including reducing model complexity using feature selection and combining relationships learned from linear algorithms into the more complex ANN architectures (i.e. a seeded ANN approach). Then, using lessons learned from predicting height, we compared the performance of all GP algorithms across all species and traits.

**Figure 1.**
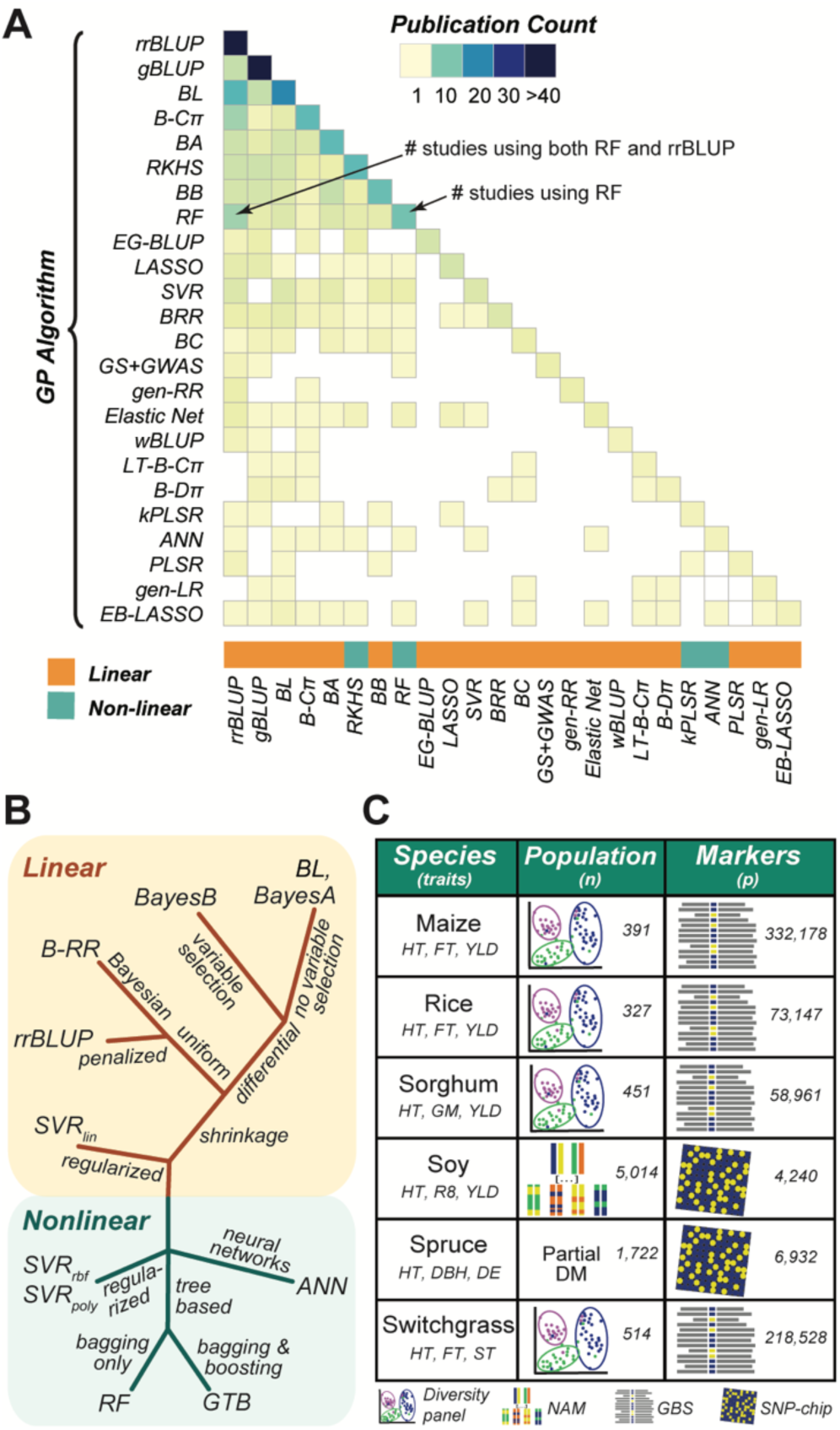
Algorithms used and compared in past GP studies and algorithms and data included in the GP benchmark. **(A)** Number of times a GP algorithm was utilized (diagonal) or directly compared to other GP algorithms (lower triangle) out of 91 publications published between 2012-2018 (**Table S6**). GP algorithms were included if they were utilized in >1 study. **(B)** Relationships and major differences between GP algorithms used in this study. rrBLUP, ridge regression Best Linear Unbiased Predictor; BRR, Bayesian Ridge Regression; BA, BayesA; BB, BayesB; BL, Bayesian LASSO; SVR, Support Vector Regression (kernel type: lin, linear; poly, polynomial; rbf, radial basis function); RF, Random Forest; GTB, Gradient Tree Boosting; ANN, Artificial Neural Network. **(C)** Species and traits included in the benchmark with training population types and sizes and marker types and numbers for each dataset. NAM: Nested Association Mapping. DM: partial diallel mating. GBS: genotyping by sequencing. SNP: single nucleotide polymorphism. HT: height. FT: flowering time. YLD: yield. GM: grain moisture. R8: time to R8 developmental stage. DBH: diameter at breast height. DE: wood density. ST: standability.

## Results

### Hyperparameter grid search is critical, particularly among non-linear algorithms

We selected six linear and five non-linear algorithms to compare their performance in GP problems (see **Methods**). While some model parameters can be estimated from the data (42), other parameters, referred to as hyperparameters, have to be user-defined (43,44). This was the case for eight of the algorithms in our study: BA, BB, SVR_lin_, SVR_poly_, SVR_rbf_, RF, GTB, and ANN. For these algorithms we conducted a grid search to evaluate the prediction accuracy of models using every possible combination of hyperparameter values (for lists of hyperparameters, see **Table S1**). To produce unbiased estimates of prediction accuracy the grid search was performed within the training set so that no data from the testing set was used to select hyperparameter values. Then we used the best set of hyperparameters from the grid search to build models using genotype and phenotype data from six plant species. This allowed us to compare the predictive performance of all algorithms included in the benchmark datasets.

To determine which hyperparameters significantly impacted model performance, we tested for changes in model performance (mean squared error; MSE) across the hyperparameter space for each algorithm/species/trait combination using Analysis of Variance (ANOVA). The degrees of freedom hyperparameter for BA and BB, both linear algorithms, that influences the shape of the prior density of marker effects (42) had no significant impact on model performance (ANOVA: *p*-value= 0.41∼1.0; **Table S2**). Other parameters for the Bayesian algorithms were determined using rules built into the BGLR package that account for factors such as phenotypic variance and the number of markers (p) (45) and were therefore not considered in our grid search. However, 15 of 16 of the hyperparameters tested for the non-linear algorithms significantly impacted performance in at least one species (**Table S2**, **Figure S1A-C**). Using height in maize as an example, we found that SVR_poly_ algorithm performed better (i.e. lower MSE) using 2^nd^ degree polynomials compared to using up to 3^rd^ degree polynomials (*p*-value = 1*10^−21^, **Figure 2A**). For RF-based models, the maximum depth (max depth) of decision trees allowed significantly impacted performance (*p*-value = 1*10^−3^, **Table S2**), with shallower trees typically performing better (**Figure 2B**). This pattern was also observed in RF models predicting height for rice, spruce, and soy (*p*-value= 1*10^−66^∼5*10^−4^, **Table S2, Figure S1B**). Because shallower decision trees are less complex, they tend not to overfit, suggesting the best hyperparameters for RF are those that reduce overfitting. The only hyperparameter from the non-linear algorithms that did not impact performance was the rate of dropout (a useful regularization technique to avoid overfitting) for ANN models, where there was no significant change in model performance when two different rates (10% and 50%) were used (*p*-value= 0.24 ∼ 0.97, **Table S2**).

**Figure 2.**
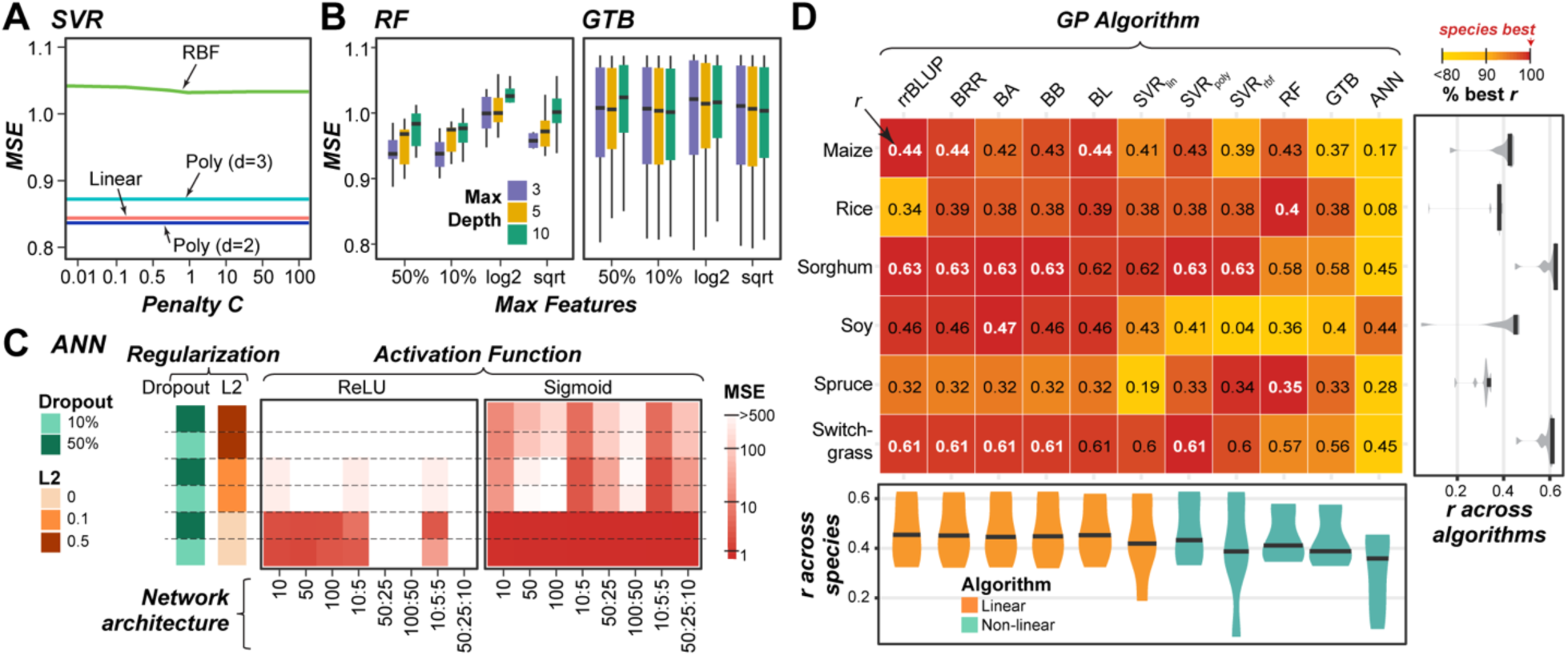
Grid search results for height in maize and overall GP algorithm performance for predicting height across species. **(A)** Average of mean squared error (MSE) over hyperparameter space (penalty, *C*) for Support Vector Regression (SVR) based models predicting height in maize. SVR_rbf_ and SVR_poly_ results are shown using gamma=1×10^−5^ and 1×10^−4^, respectively. Poly: polynomial. RBF: Radial Basis Function. **(B)** Distribution of MSEs across hyperparameter space for Random Forest (RF; left) and Gradient Tree Boosting (GTB; right) as the maximum features available to each tree (Max Features) and maximum tree depth (color) change. GTB results are shown using a learning rate = 0.01. **(C)** Average MSE across hyperparameter space for ANN models with different network architectures, degrees of regularization (dropout or L2), using either the Rectified Linear Unit (ReLU; left) or Sigmoid (right) activation function. **(D)** Mean performance (Pearson’s Correlation Coefficient: *r*, text) for predicting height and percent best *r* (colored box, top algorithm for each species = 100% (red)). White text: the best *r* values. Violin-plots show the median and distribution of *r* values for each trait (right) and algorithm (bottom).

### ANN is the most significantly impacted by hyperparameter choice

Hyperparameters for SVR_lin_, SVR_poly_, SVR_rbf_, RF, and GTB tended to have moderate effects on MSE, while ANN hyperparameters often caused substantial changes in MSE (**Figure 2A-C; Figure S1A-C**). Across the six species, the median variance in MSE across the hyperparameter space for ANN was 6*10^6^, but ranged from 3*10^−3^ – 0.1 for the other GP algorithms (**Figure S1D**) For example, for predicting height in maize, SVR_poly_ models built using the 2^nd^ degree polynomial outperformed those built using the 3^rd^ degree polynomial with a decrease in MSE ∼ 0.05 (**Figure 2A**), while for ANN models, hyperparameter combinations that performed the best (i.e. Sigmoid activation function and no L2 regularization) resulted in models with MSEs that were >500 lower than the worst performing model (Rectified Linear Unit (ReLU) activation function, no L2 regularization, and large numbers of hidden nodes; **Figure 2C**). This highlighted that, while hyperparameter selection is necessary for all non-linear algorithms, it is especially critical for building ANNs for GP problems.

Using the best set of hyperparameters for each model, we next compared the predictive performance (Pearson’s correlation coefficient, *r*, between predicted and true trait values) of each algorithm on plant height. As with past efforts to benchmark GP algorithms (23,24), no one algorithm always performed the best (white bolded; **Figure 2D**). For example, while rrBLUP performed best for maize, sorghum, and switchgrass, BA performed best for soy, and RF performed best for rice and spruce. Notably, ANNs substantially underperformed compared to other non-linear algorithms, with a median performance at 84% of the best *r* for each of the six species (i.e. 16% below the best performing algorithm for that trait/species).

Notably, among the six species, ANN performed the best in soy (*r* = 0.44) relative to the species best algorithm BA (*r* = 0.47, **Figure 2D**). Soy has the largest number of training lines among the six species (5,014) and has a marker to training line ratio close to one (**Figure 1C**). Thus, we hypothesized the poor performance of the ANN models was in part due to our inability to train a network with so many features (markers) and so little training data (lines). During ANN model training, the weights assigned to each connection between nodes in neighboring layers of the network have to be estimated. Because every input marker is connected to every node in the first hidden layer, including more markers in the model will require more weights to be estimated, resulting in a more complex network that is more likely to underfit. In an ideal situation, to account for the complexity in these large networks, five to ten times more instances (lines) than features (markers) would need to be available for training (46). Alternatively, one can reduce model complexity by only including markers that are most likely to be associated with the trait using feature selection methods.

### Feature selection improves performance of ANN models

ANNs and sometimes other non-linear algorithms performed poorly compared to linear methods, which could be due to an insufficient number of training lines relative to the number of markers. To address this, we used feature selection to identify and select the markers most associated with trait variation. Because the number of markers associated with a trait is dependent on the genetic architecture of the trait and is not typically known, models were built using a range of numbers of markers (p = 10∼8,000) and were compared to models built using all available markers from each species. Because performing feature selection on the training and testing data can artificially inflate prediction accuracies (47), feature selection was conducted on the training set only. This was repeated 10 times, using a different subset of lines for testing for each replicate (**see Methods**).

Three feature selection algorithms (RF, BayesA, and Elastic Net (EN)) were compared to predict height in maize, the species with the largest number of markers (p) relative to training lines (n) (p:n = 850, **Figure 1C**). While each algorithm selected a largely different subset of markers (**Figure 3A**, **Figure S2A**), the degree of overlap was significantly greater than random expectation. To demonstrate this, we randomly selected three sets of 8,000 maize markers and counted how many markers were present in all three sets 10,000 times and found that the 99^th^ percentile of overlap was equal to 10, however we observed an average of 220 overlapping markers across replicates using these three feature selection approaches. When the different feature selection subsets were used to predict height in maize, there was a significant interaction between the number of available markers (p) and the feature selection method (ANOVA: *p*- value = 3*10^−7^). Exploring this interaction further, we found that, while feature selection algorithms performed similarly with large n, RF tended to perform the best when fewer markers were selected for GP (**Figure 3B; Figure S2B**) and was therefore used to test the impact of feature selection on predicting height in the other five species.

**Figure 3.**
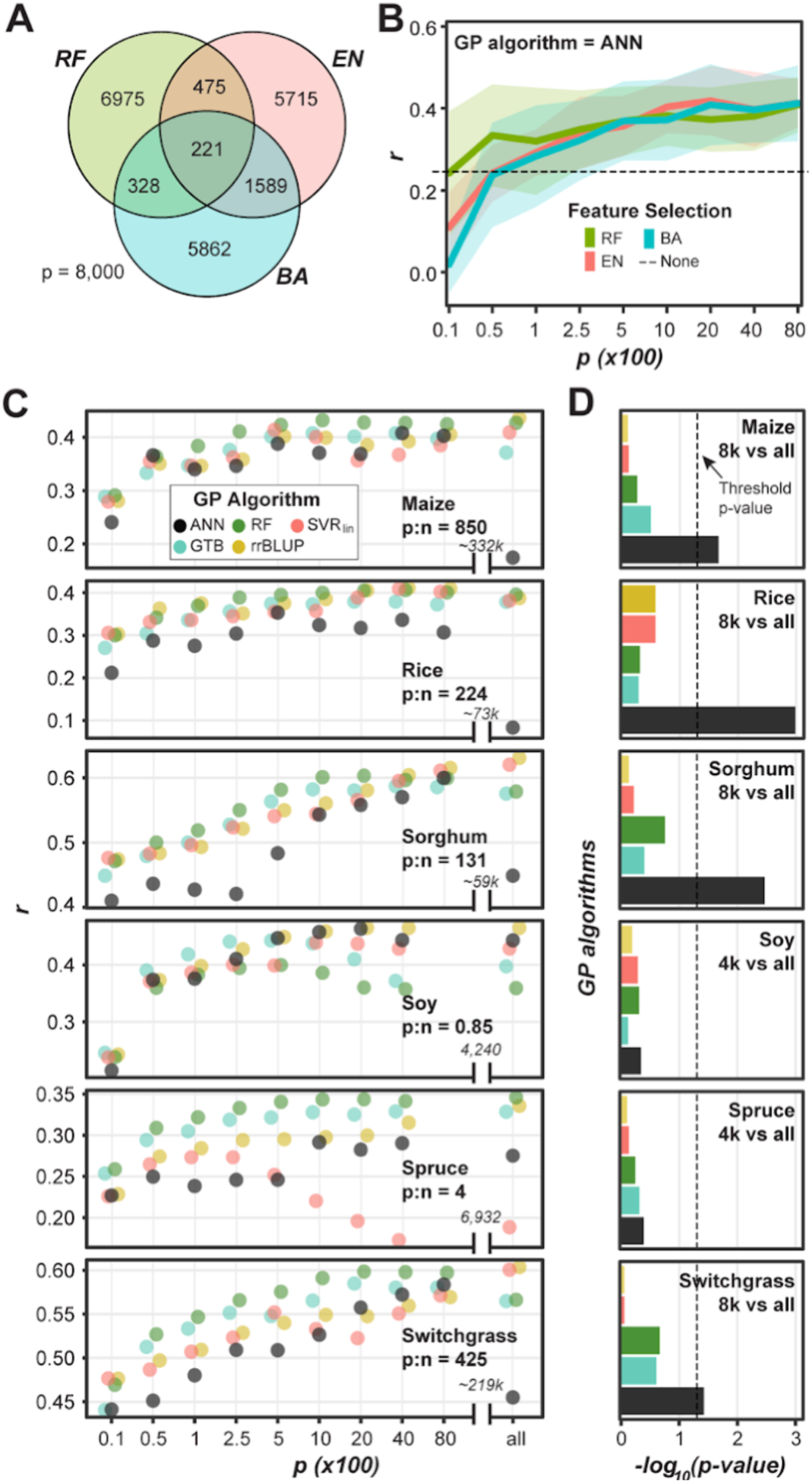
Impact of feature selection on GP algorithm performance. **(A)** Average number of overlapping markers in the top 8,000 markers selected by three feature selection algorithms for predicting height in maize across ten replicates. EN: Elastic Net. **(B)** Change in ANN predictive performance (*r*) at predicting height in maize as the number of input markers (p) selected by three feature selection algorithms (BayesA: BA, EN, and Random Forest: RF) increases. Dashed line: mean *r* when all 332,178 maize markers were used. **(C)** Mean *r* of rrBLUP, SVR_lin_, RF, GTB, and ANN models for predicting height using subsets or all (X-axis) markers as features across 10 replicate feature selection and ML runs for each of six species with their ratios of numbers of markers (p) to numbers of lines (n) shown. Data points were jittered horizontally for ease of visualization. **(D)** The significance (-log_10_(*p*-value), Mann-Whitney U test) of the difference in *r* between models from different GP algorithms (colored as in **Figure 3C**) generated using a subset of 4,000 or 8,000 and all markers as input. Dotted line designates significant differences (*p-*value < 0.05).

For species with a low p:n ratio (i.e. soy and spruce), for all GP algorithms tested, as p increased the model performance tended to increase continuously (e.g. all GP algorithms in sorghum) or, in some cases, the model performance reached a maximum (or a plateau) quickly (e.g. in soy after 2,500 markers were used) (**Figure 3C**). We used a one-sided Mann Whitney U (MWU) test to determine if models using a selected subset (p = 4,000) of markers performed better than models using all markers. For these species, we found no significant improvement in performance after feature selection using any GP algorithm (*p*-value = 0.41 ∼ 0.79; **Figure 3D**). For example, ANNs built using all 6,932 spruce markers performed no better than those built using the top 4,000 markers (*p*-value= 0.41).

For species with a large p:n ratio (i.e. maize, rice, sorghum, and switchgrass), a similar pattern was observed for rrBLUP, SVR_lin_, RF, and GTB, where performance increased or reached a plateau as p increased and no significant improvement in performance was found after feature selection (p=8,000) (*p*-value= 0.18 ∼ 0.88; **Figure 3D**). However, for these four species, feature selection dramatically improved the performance of ANN models (*p*-value= 0.001 ∼ 0.038; **Figure 3D**). For example, after feature selection prediction of height in maize using ANNs improved from *r*=0.17 to 0.41, a 141% increase. Ultimately, performing feature selection prior to ANN training for these four datasets with large p:n ratios, improved ANN performance (median *r* at 89% of the best *r* for each of the six species) compared to ANNs without feature selection (84% of the best *r*). Therefore, for the GP benchmark analysis, feature selection was performed prior to model building for additional traits for maize, rice, sorghum, and switchgrass and the top 8,000 markers were used.

While feature selection notably improved ANN performance, ANNs still often underperformed compared to other GP algorithms (**Figure 3C**), meaning the they were unable to learn even the linear relationships between markers and traits that were found using the linear-based algorithms. Because ANNs should theoretically at least match the performance of linear algorithms, this suggests that the ANN hyperparameters are not optimal. Furthermore, we found that, even after feature selection, there was greater variation in performance across replicates for ANN models compared to rrBLUP, SVM_lin_, RF, and GB (**Figure S2C-D**), indicating the ANN models did not always converge on the best solution. One potential reason for the is that the final trained network can be heavily influenced by the initial weights used in ANN, which are selected randomly. In addition, while random weight initialization, a procedure we have used thus far, reduces bias in the network, it can also result in some networks converging on a local, rather than global, optimal solution.

### Non-random initialization of ANN starting weights improves ANN performance for some species

To reduce the likelihood of ANNs converging to locally optimal solutions, we developed an approach that allowed the ANNs to utilize the relationship between markers and traits determined by another GP algorithm. In this approach, a GP algorithm was applied to the training lines, and the coefficient or importance score assigned to each marker from this algorithm was used to seed the starting weights (**Figure 4A**). Four GP algorithms were tested to seed the weights: rrBLUP, BB, BL, and RF (referred to as ANN_rrBLUP_, ANN_BB_, ANN_BL_, and ANN_RF_, respectively). Because this approach could predispose the networks to only learn the relationship already identified by the seed algorithm, two steps were taken to re-introduce randomness into the network (see **Methods**). First, the seeded approach was only used to initialize starting weights for 25% of the nodes in the first hidden layer, while connection weights to the remaining 75% of nodes were initialized randomly as before. Second, noise was infused into the starting weights for the 25% of nodes that were seeded.

**Figure 4.**
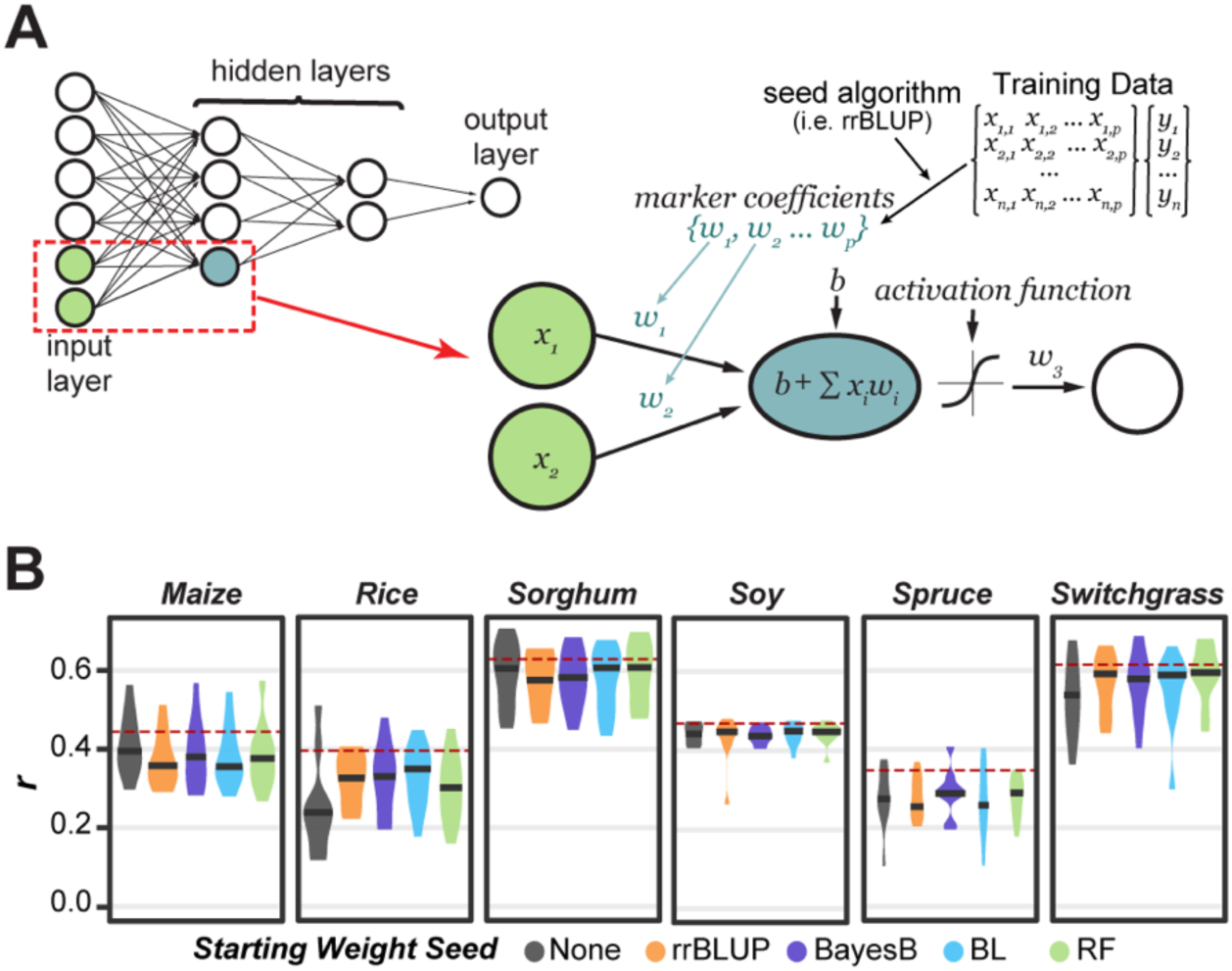
Description and performance results of the seeded ANN Approach. **(A)** An overview of the seeded ANN approach. The network in the top left is an example of a fully connected ANN with 6 input nodes (i.e. 6 markers), two hidden layers, and one output layer (i.e. predicted trait value). The blue node in the first hidden layer represents an example node that will have seeded weights. For this node, the weights (*w*) connecting each input node to the hidden node will be seeded from the coefficient/importance for each marker as determined by another GP algorithm using the training data. *b*: bias, which helps control the value at which the activation function will trigger. **(B)** The distribution of model performance (*r*) using only all random (None) or 25% seeded (rrBLUP, BayesB, BL, RF) weight initialization. The mean performance of the overall top performing algorithm (i.e. not necessary ANN) shown as dotted red line.

Applying this approach to predict plant height we found that ANN performance improved for three of six species (**Figure 4B**). For example, the median performance for rice without seeding (ANN) was *r* = 0.24 and with seeding from BL (ANN_BL_) was *r* = 0.35, a 46% improvement, while for sorghum, ANN_BL_ had <0.1% improvement over the original ANN methods. Seeding ANN models did not significantly reduce the amount of variation in model performance across replicates (ANOVA: *p*-value= 0.81, **Table S5**). Ultimately, seeded ANN models had a median performance between 89% - 90% of the best *r* for each species (compared to 89% with random initialization, **Figure 4B**). While this represented only a moderate improvement, we included the seeded ANN approach in the benchmark analysis because of how substantial the improvement was for some species (i.e. rice).

### No one GP algorithm performs best for all species and traits

Having established best practices for hyperparameter and feature selection for our datasets, we next compared the performance of all GP algorithms for predicting three traits in each of the six species. For maize, rice, and soy, these traits included height, flowering time, and yield (**Figure 1C**). For species where data was not available for one or more of these traits, other traits were used (see the panel labeled “Others”, **Figure 5A**). As with past efforts to benchmark GP algorithms (23,24), different algorithms performed best for different species/trait combinations (**Figure 5A**; **Table S4**). Thus, we utilized the predictive power of multiple algorithms to establish an ensemble prediction using all (EN_11_) or a subset of five (EN_5_) algorithms (see **Methods**). The ensemble models consistently performed well, with EN_5_ or EN_11_ being the best (three) or tied for the best (nine) algorithm for 12 of thse 18 species/trait combinations included in the benchmark and had a median performance rank of 2.5 (**Figure 5B; Table S5**). For the remaining 6 species/trait combinations where EN_5_ or EN_11_ weren’t among the best performers, they tended to perform only slightly worse (median % of best *r* = 97.2%, **Figure 5A**). Taken together this suggests that ensemble-based predictions are more stable and more likely to result in better trait predictions.

**Figure 5.**
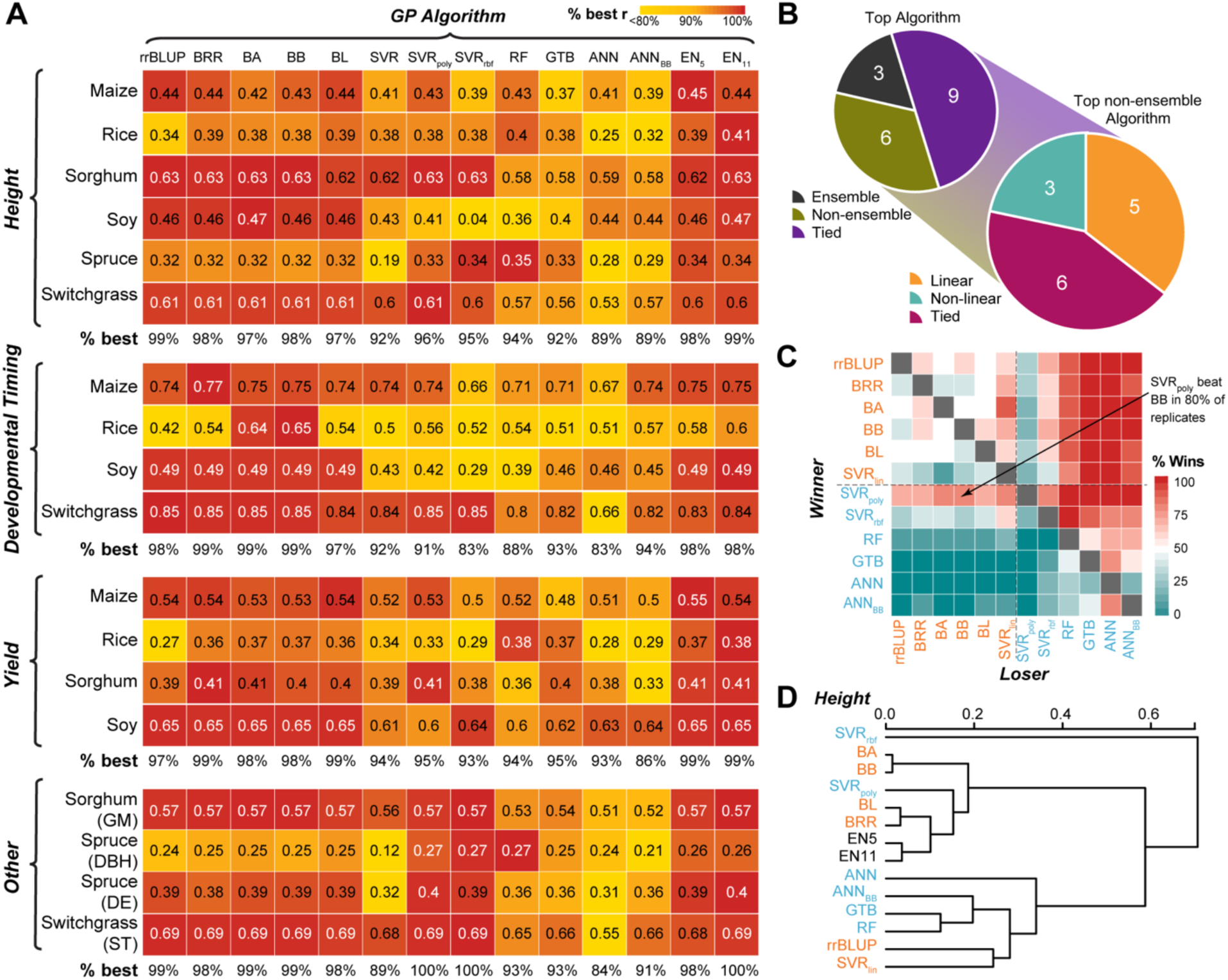
Comparison of algorithms for predicting additional traits. **(A)** Mean model performance (*r*; text) for each species/trait combination (y-axis) for each GP algorithm (x- axis). White text: *r* of the best performing algorithm(s) for a species. Colored boxes: percent of best performance (*r*) for a species, with the top algorithm for each species = 100% (red). The median % of best performance for each GP algorithm for each type of trait (i.e. height, developmental timing, yield, other) is shown below each heatmap. GM: sorghum grain moisture. DBH and DE: diameter at breast height and wood density, respectively, for spruce. ST: standability for switchgrass. **(B)** Top left: summary of the number of species/trait combinations that were predicted best by an ensemble (gray) or a non-ensemble model (yellow), or predicted equally well by both (purple). Bottom right: among non-ensemble models that performed or tied for the best, the number of species/trait combinations that were predicted best by a linear (blue) or a non-linear model (green) or predicted equally well by both (orange). **(C)** Percent of replicates where one GP algorithm (y- axis, winner) outperformed another GP algorithm (x-axis, loser) for predicting height in switchgrass. Orange and cyan texts: linear and non-linear algorithms, respectively. **(D)** Hierarchical clustering of GP algorithms based on mean predictive performance across all species/trait combinations. Algorithm colored as in **(C)**.

Focusing on the species/trait combinations where one of the non-ensemble algorithms was or tied for best, we found that a linear algorithm performed best for five of the species/trait combinations, a non-linear algorithm performed best for three species/trait combinations, and both a linear and a non-linear algorithm performed equally well for the remaining six species/trait combinations (**Figure 5B**). This finding suggests that linear and non-linear algorithms are equally well suited for GP. The linear algorithms BA and BRR performed best overall, being among the top performers for 9 traits and with the top two median ranks of four and five, respectively (**Table S5**). The top performing non-linear algorithm was SVR_poly_, which was among the top performers for 8 traits and had a median rank of 6. There was notably greater performance variation across species/traits for non-linear algorithms (mean variance = 1.16%) compared linear algorithms (mean variance = 0.32%) (**Table S5**). For example, SVR_rbf_ performed poorly at predicting developmental timing traits (median 83% of the best *r*), however it had or was tied for the best prediction for three of the four “other” traits (median 100% of the best *r*) (**Figure 5A**). Results from ANN models using randomly initialized (ANN) and BB seeded (ANN_BB_) weights are shown because ANN_BB_ had the best performance of the seeded ANN models (see Supplemental for results from other seeded ANNs). Notably, neither the randomly initialized ANN (median rank = 13.5) nor the ANN_BB_ (median rank = 13) models performed best for any trait (**Table S5**).

One limitation of comparing the mean score or performance rank is that small but consistent differences in model performance could be missed. To account for this, we also calculated the number of times an algorithm outperformed another algorithm for each trait across the replicates. Using this metric, we were able to identify algorithms that consistently outperformed others for a given trait/species combination (**Figure 5C, Figure S3**). We frequently observed that linear algorithms had higher win percentages than nonlinear algorithms, this was the case for all three traits in maize and soybean for example (**Figure S3**). However, there were plenty of exceptions. RF and SVR_rbf_ had higher win percentages than linear algorithms for predicting height and diameter at breast height (DBH) in spruce and ANN_BB_ had a higher win percentage than all algorithms except BA and BB for predicting flowering time in rice (**Figure S3**). In a few cases, assessing win percentages allowed us to identify winners when mean predictive performance (*r*) was tied. For example, for predicting height in switchgrass. SVR_poly_ had the same average performance (*r* = 0.61) as multiple of the linear algorithms (i.e. rrBLUP, BA, etc.), however, it outperformed those algorithms in 70-80% of replicates (**Figure 5C**).

Finally, we performed hierarchical clustering of the algorithms based on their performance across the 18 species/trait combinations (from **Figure 5A**) in order to group similarly predictive algorithms. Interestingly, linear and non-linear algorithms did not clearly separate from each other (**Figure 5D**). For example, rrBLUP and SVR_lin_ were more similar to the non-linear, decision tree-based algorithms (i.e. RF and GTB) than they were to the linear Bayesian algorithms (i.e. BA and BRR). Notably, while the Bayesian algorithms tended to cluster together closely, the non-linear algorithms were often highly different, with SVR_rbf_ and ANN algorithms performing most uniquely.

## Discussion

We conducted a benchmarking comparison of GP algorithms on 18 species/trait combinations that differ in the type and size of the training data set and of the marker data available. A key result from this analysis is that no one model performs best for all species and all traits. With that said, linear algorithms tend to perform consistently well, while the performance of non-linear algorithms varied widely by trait. Studies of gene networks have shown that non-additive interactions (e.g. epistasis, dominance) are important for development and regulation of complex traits (13,14). One may expect approaches that can consider non-linear combinations would therefore be better suited for modeling complex trait. This was not the case and we found the inconsistency of non-linear algorithms surprising.

We have three, non-mutually exclusive, explanations for why linear algorithms often outperform non-linear algorithms. First, the traits included in this study vary in their genetic architecture (i.e. the number and distribution of allele effects), therefore we may be observing that linear algorithms outperform non-linear algorithms when the trait has a predominantly additive genetic basis. Second, there is evidence that even highly complex biological systems generate allelic patterns that are consistent with a linear, additive genetic model because of the discrete nature of DNA variation and the fact that many markers have extreme allele frequencies. In diploid organisms, at each locus there are three possible variants (e.g. AA, TT, AT). However, when allele frequencies are extreme, most of the genotypes fall within two categories. When this happens, a linear model can capture (almost) all the variance generated by each locus, even those under dominant gene action. This phenomena also affects the proportion of epistatic variance that can be captured by additive model (48), which can explain why additive models often perform very well at predicting traits that at the biological level are affected by complex epistatic networks. Finally, a third explanation is that the amount of training data available for most GP problems was insufficient for learning non-linear interactions between large numbers of markers, therefore the linear models, which focus on modeling linear relationships, outperform the non-linear models.

A number of findings from our study indicate that limited training data plays a role. We found that non-linear algorithms (i.e. RF, SVR_poly_, and SVR_rbf_) performed best at predicting traits in spruce, the species with the second smallest marker density to training population size ratio. In addition, ANN models tended to perform better at predicting traits in soy, the species with the lowest marker density relative to the training data set size (p:n) included in this study. However, a recent study involving very large sample size (n∼80,000) in humans compared linear models with two types of ANN algorithms, multilayer perceptron and convolutional neural networks, and did not find any clear superiority of the ANN methods relative to linear models, if anything the linear model offered higher predictive power than the ANNs (37).

Furthermore, the ANN models significantly improved after feature selection. This was not the case for other algorithms in our study or with previous efforts to use feature selection for GP (47,49). For example, for predicting traits in Holstein cattle, the top 2,000 markers had only 95% of the predictive ability of all the markers using BL (49). With a fixed training data size, prediction accuracy is a function of how much genetic variation is captured by markers in linkage disequilibrium with quantitative trait loci and the accuracy of the estimated effects (50). Because feature selection removes markers from the model, such decreases in performance after feature selection for non-ANN models are likely due to the reduction in the amount of genetic variation captured without a subsequent increase in the accuracy of the estimated effects. However, we hypothesize that feature selection significantly improved performance for ANNs because it improved the accuracy of the estimated effects (i.e. the connection weights) more than it reduced the amount of genetic variation captured. A similar observation was made by Bellot *et al.*, as they found that ANN models using the top 50k markers performed no better, and in some cases performed worse, than ANN models using the top 10k markers (37).

While there is a great deal of excitement about the uses of deep learning in the field of genetics, there is still much work to be done to improve performance of deep learning-based models. In this study we identified dimensionality as a major limitation to training ANNs for GP. Additional areas of deep learning research also need to be further explored. For example, in this study we limited the hyperparameter space searched because the grid search method was too computationally intensive to be more thorough. Because changes in hyperparameters had a large impact on model performance, further hyperparameter tuning could lead to better performing models. For example, we limited our search to include nine possible network architectures with between one and three hidden layers each containing between 5-100 nodes (**Table S1**), but it is possible that ANNs with different network architectures, such as more hidden layers, or different combinations of layer sizes, could have performed better. These caveats aside, we found that the ANNs performed better when a large number of training examples (n) were available. This suggests that these models may be useful for larger breeding programs.

## Conclusion

In summary, we provided a thorough comparison of 11 GP algorithms for predicting diverse traits in six plant species with a range of marker types and numbers and population types and sizes. We found that the performance of ensemble models, generated by combining predictions from multiple individual GP algorithms, consistently tied with or exceeded the performance of the best individual algorithm. Because of this and our finding that no GP algorithm was best for all species/trait combinations, we recommend that breeders test the performance of multiple algorithms on their training population to identify which algorithm or combination of algorithms performs best for traits important to their breeding program. While for some species and traits the performance difference between algorithms may be small, over the course of an entire breeding program, these small differences could add up to large genetic gains.

## Methods

### Genotype and phenotype data

Genotypic data from six plant species were used to predict 3 traits from each species (20,51–53) (**Fig 1C**). The maize phenotypic (54) and genotypic (55) data were from the pan-genome population. The rice data were from elite breeding lines from the International Rice Research Institute irrigated rice breeding program (20), and dry season trait data averaged over four years were used. The sorghum data were generated from sorghum lines from the US National Plant Germplasm System grown in Urbana, IL (51) and trait values were averaged over two blocks for this study. The soybean data were generated from the SoyNAM population containing recombinant inbred lines (RILs) derived from 40 biparental populations (52). The white spruce data were obtained from the SmartForests project team, using a SNP-chip developed by Quebec Ministry of Forest Wildlife and Parks (^56^). Switchgrass phenotypic (53) and genotypic (57) data were generated from the Northern Switchgrass Association Panel (58) which contains clones or genotypes from 66 diverse upland switchgrass populations.

The genotype data was obtained in the form of biallelic SNPs with missing marker data already dropped or imputed by the original authors. Marker calls were converted when necessary to [-1,0,1] corresponding to [aa, Aa, AA] where A was either the reference or the most common allele. Genome locations of maize SNPs were converted from assembly AGPv2 to AGPv4, with AGPv2 SNPs that did not map to AGPv4 being removed, leaving 332,178 markers for the maize analysis. Phenotype values were normalized between 0 and 1 and, when necessary, were averaged over replicates, blocks, locations, and/or years. Lines with missing phenotypic value for any of the three traits were removed.

### Genomic selection algorithms

To assess what statistical approaches are most frequently used for genomic selection, we conducted a literature search of papers applying genomic selection methods to crop or simulated data from January 2012-February 2018. We recorded what statistical approach(es) was(were) applied in each study (**Table S6**), allowing us to calculate both the total number of times an approach had been applied and how many times any two approaches were directly compared (**Fig 1A**). Based on the results from this literature search, nine commonly used statistical approaches were included in this study: rrBLUP, Bayes A (BA), Bayes B (BB), Bayesian LASSO, Bayesian-RR, RF, SVR with a linear kernel (SVR_lin_), SVR with polynomial kernel (SVR_poly_), SVR with radial basis function kernel (SVR_rbf_). Two additional machine learning approaches, gradient tree boosting (GTB) and artificial neural networks (ANN), were also included because of their ability to model non-linear relationships.

Most linear algorithms were implemented in R packages rrBLUP(59) and BGLR (for Bayesian methods including BRR: Bayesian RR, BA: Bayes A, BB: Bayes B, and BL: Bayesian LASSO)(45). These algorithms vary in what approach they use to address the p >> n problem (**Figure 1B**), for example rrBLUP performs uniform shrinkage on all marker coefficients to reduce variance of the estimator, while BB performs differential shrinkage of the marker coefficients and variable selection. The differences between these algorithms have been thoroughly reviewed previously (42). Models for Bayesian methods were trained for 12,000 iterations using a burn-in of 2,000.

Non-linear algorithms (SVR_poly_, SVR_rbf_, RF, and GTB) and SVR_lin_ were implemented in python using the Scikit-Learn library (60). For SVR algorithms, the marker data is mapped into a new feature space using linear or non-linear kernels (i.e. poly, rbf) and then linear regression within that feature space is performed with the goal of minimizing error outside of a margin of tolerated error. The RF algorithm works by averaging the predictions from a “forest” of bootstrapped regression trees, where each tree contains a random subset of the lines and of the markers (61). Related to RF, GTB algorithm uses the principle of boosting (62) to improve predictions from weak learners (i.e. regression trees) by iteratively updating the learners to minimize a loss function, therefore generating better weak learners as training progresses.

Artificial Neural Networks (ANNs) were implemented in python using TensorFlow (63). The input layer for the ANNs contained the genetic markers for an individual (*x*; **Figure 1B**), the nodes in the hidden layers were all fully connected to all nodes in the previous and following layers (i.e. Multilayer Perceptron). A non-linear activation function (selected during the grid search, see below) was applied to each node in the input and hidden layers, except the last hidden layer, which was connected with a linear function to the output layer, the predicted trait value (*y*). To reduce the likelihood of vanishing gradients, when the error gradient, which controls the degree to which the weights are updated during each iteration of training, becomes so small the weights stop updating thus halting model training, in the ANN, the starting weights (*w*) were scaled relative to the number of input markers using the Xavier Initializer (64). Weights were then optimized using the Adam Optimizer (65) with a learning rate selected by the grid search (described below). To determine the optimal stopping time for training (i.e. number of epochs), an early stopping approach was used (66), where the training set was further divided into training and validation, and early stopping occurred when the change in mean squared error (MSE) for the validation set was < 0.1% for 10 epochs using a 10 epoch burn-in. Occasionally, due to poor random initialization of weights, the early stopping criteria would be reached before the network started to converge and the resulting network would predict the same trait value for every line. When this was observed in the validation set the training process was repeated starting with new initialized weights.

To incorporate predictions from multiple algorithms into one summary prediction, an ensemble approach was used where the ensemble predicted trait value was the mean predicted trait value from all algorithms (EN_11_) or a subset of algorithms (EN_5_: rrBLUP, BL, SVMpoly, RF, ANN) algorithms. The subset consisted of algorithms with differing statistical bases, where rrBLUP represented penalized methods, BL represented the Bayesian approaches, SVMpoly represented non-linear regularized functions, RF represented decision tree based methods, and ANN represented the deep learning approach. This ensemble predicted trait value was then compared to the true trait values to generate performance metrics.

### Hyperparameter grid search using cross-validation

To obtain the best possible results from each algorithm, a grid search approach was used to determine the combination of hyperparameters that maximized performance for each trait/species combination. No hyperparameter needed to be defined for rrBLUP, BL, or BRR. For rrBLUP, the R package estimates the regularization and kernel parameters from the data. For BL or BRR, parameters for these Bayesian regression methods were also estimated from the data. Between one and five hyperparameters were tested for the remaining algorithms (**Table S3**).

To avoid biasing our hyperparameter selection, an 80/20 training/testing approach was used, where 20% of the lines were held out from each model as a testing set and the grid search was performed on the remaining 80% of training lines. For RF, SVR_lin_, SVR_poly_, SVR_rbf_, and GTB algorithms, 10 replicates of the grid search were run using the GridSearchCV function from Scikit-Learn with 5-fold cross validation. Ten replicates of the grid search were also run for ANN models, where for each replicate 80% of the training data was randomly selected for training the network with each combination of hyperparameters and the remaining 20% used to select the best combination. This whole process (train/test split, grid search) was replicated 10 times, with a different 20% of lines selected as the test set for each replicate. Analysis of variance (ANOVA) implemented in R was used to determine which hyperparameters significantly impacted model performance for each species.

### Assessing Predictive Performance

The predictive performance of the models was compared using two metrics. For the grid search analysis, the mean squared error (MSE) between the predicted (Ŷ) and the true (Y) trait value was used. For the model comparisons, Pearson correlation coefficient (*r*) between the predicted (Ŷ) and the true trait value (Y) was used as it is the standard metric for GP performance (2,24,29). It was computed using the cor() function in R for rrBLUP and the Bayesian approaches or the numpy corrcoef() function in Python for the ML and ANN approaches. Only predicted trait values for lines from the test set were considered when calculating *r*. Summary performance metrics (% of best *r,* rank, variance) were calculated using the mean predictive performance (*r*) across all replicates for each GP algorithm for each species/trait combination.

### Feature Selection

The top 10, 50, 100, 250, 500, 1000, 2000, 4000, and 8000 markers were selected using three different feature selection algorithms: Random Forest (RF), Elastic Net (EN), and BayesA (BA). RF and EN feature selection were implemented in Scikit-Learn and BA was implemented in the BGLR package in R. The EN feature selection algorithm requires tuning of the hyperparameter that controls the ratio of the L1- and L2- penalties (e.g. L1:L2 = 1:10 = 0.1). Because the L1 penalty function performs variable selection by shrinking some coefficients to zero, we started with an initial weight on the L1 penalty of 0.1 and then, if fewer than 8,000 markers remained after variable selection, we reduced it in steps of 0.02 until that criteria was met (a 4,000 marker threshold was used for spruce and soy, which only had 6,932 and 4,240 markers available, respectively).

To avoid bias during feature selection, the 80:20 training/testing approach described above was used, where feature selection was performed on the training data and the ultimate performance of models built using the selected markers was scored on the testing set.

### Initializing ANN starting weights seeded from other GP algorithms

In addition to building ANNs with randomly initialized starting weights, we tested the usefulness of seeding the starting weights with information from other GP algorithms (i.e. rrBLUP, BB, BL, or RF) (**Figure 4A**). This is an ensemble-like approach in that it utilizes multiple algorithms to make a final prediction. Ensemble approaches often perform better than single algorithm approaches (67). First, after the data was divided into training, validation, and testing sets and, for species with large p:n ratios (i.e. maize, rice, sorghum, switchgrass) the top 8,000 markers were selected, we applied a GP algorithm (rrBLUP, BB, BL, or RF) to the training data. From that model we extracted the coefficients/importance scores assigned to each marker and used those as the starting weights for 25% of the nodes in the first hidden layer. We also tested seeding starting weights for 50% of the nodes to predict height in all 6 species but found this significantly increased the model error (MSE) on the validation set (ANOVA; *p*- value= 0.04), so only results from seeding 25% were included. Because we still needed to reduce the likelihood of vanishing gradients, described above, we manually adjusted the scale of the coefficients/importance scores to match the distribution of the starting weights assigned the remaining 75% of the nodes in the first hidden layer by Xavier Initialization. Finally, to reduce bias in the ANN, random noise was introduced to the seeded nodes by multiplying each starting weight with a random number from a normal distribution with a mean =0 and the standard deviation equal to the standard deviation of weights from Xavier Initialization.

After the training data was used to determine these seeded starting weights, it was used to train the ANN model, the validation set was used to select the best set of hyperparameters and the early stopping point. Then the final trained model was applied to the testing set and performance metrics were calculated.

### Data and Code Availability

The datasets supporting the conclusions of this article will be made publicly available on Dryad at the time of publications. The scripts to run all of the algorithms included in this study are available on GitHub. The R code for all statistical analyses included in the manuscript is available at (https://github.com/ShiuLab/Manuscripts/2019_GP_comparison). Code for running rrBLUP and Bayesian algorithms is available at (https://github.com/ShiuLab/GenomicSelection). The pipeline for running SVR, RF, GTB (i.e. machine learning algorithms) is available at (https://github.com/ShiuLab/ML-Pipeline). This pipeline also includes code for performing feature selection (https://github.com/ShiuLab/ML-Pipeline/FeatureSelection.py). The pipeline for running ANN (i.e. deep learning algorithms) is available at (https://github.com/ShiuLab/ANN_Pipeline). This pipeline allows for the user to select randomly initialized starting weight (default) or seeded starting weights (e.g.: -weights BayesB).

## Author Contributions

CBA, GC, and S-HS conceptualized the study, CBA, AM, MR, and GC developed the methodology. Data curation, formal analysis, software development, visualizations, and writing the original draft was done by CBA. All author read, edited, and approved the final manuscript.

## Abbreviations

GP: genomic prediction
ANN: artificial neural network
rrBLUP: ridge regression best linear unbiased prediction
BA: Bayes A
BB: Bayes B
LASSO: least absolute angle and selection operator
BL: Bayesian LASSO
SVR: support vector regression
lin: linear
rbf: radial basis function
poly: polynomial
RF: random forest
GTB: gradient tree boosting
p: number of markers
n: number of lines
MSE: mean squared error
ANOVA: analysis of variance
ReLU: rectified linear unit
EN: elastic net
MWU: Mann Whitney U
DBH: diameter at breast height

## Acknowledgements

We thank Peipei Wang and John Lloyd from the Shiu lab, Gabriel Rovere from the MSU QuantGen group, and Fouad Bahrpeyma from the Insight Center for their valuable suggestions to our project. This work was partly supported by the National Science Foundation Graduate Research Fellowship (Fellow ID: 2015196719), Graduate Research Opportunities Abroad (GROW) Fellowship to C.B.A.; the U.S. Department of Energy Great Lakes Bioenergy Research Center (BER DE-SC0018409) and National Science Foundation (IOS-1546617, DEB-1655386) to S.-H.S; and the National Institute of Health (R01GM099992, R01FM101219) to G.D.L.C.. The funding bodies had no role in the design, analysis, interpretations, or writing of this study.

## Supplemental Legends

**Figure S1. Height prediction performance for non-linear GP algorithms during hyperparameter grid search**.

**(A)** Average (line) and standard deviation (shadow) of mean squared error (MSE) over hyperparameter space for SVR based models predicting height as the penalty (*C*) (X-axis) change. SVR_rbf_ and SVR_poly_ results are shown using gamma=1×10^−5^ and 1×10^−4^, respectively. **(B)** Distribution of the MSE across hyperparameter space for RF (left) and GTB (right) as the maximum features available to each tree (Max Features; X-axis) and maximum tree depth (color) change. GB results are shown using a learning rate = 0.01. **(C)** Average MSE across hyperparameter space for ANN models with different network architectures (X-axis), degrees of regularization using dropout (D.o.) or L2 regularization (L2), using either the Rectified Linear Unit (ReLU; left) or Sigmoid (right) activation function. **(D)** Distribution of the variance in MSE across the hyperparameter space for predicting height in each species using each GP algorithm. Black bar represents the median variance across the species for each GP algorithm.

**Figure S2. Comparison of feature selection algorithms and change in performance variation after feature selection**.

**(A)** Average number of overlapping markers in the top markers (p) selected by three different feature selection algorithms for predicting height in maize across ten replicates for p=10 ∼ 8,000. **(B)** Change in model performance (*r*) using five GP algorithms at predicting height in maize as the number of input markers (p) selected by three different feature selection algorithms increases. Dashed line: the mean *r* for each GP algorithm when all maize markers were used. Colored lines: mean *r* of models using features selection subsets using algorithms colored as in **(A)**. Colored areas: standard deviation around the mean. **(C)** Distribution and median of the standard deviation of model performance (*r*) across replicates for all feature selection subsets (p=10 ∼ 8,000) combined across all species for each GP algorithm **(D)** Distribution and median of the standard deviation of model performance across replicates for all feature selection subsets (p=10 ∼ 8,000) by species for each GP algorithm.

**Figure S3. Number of wins between each pair of GP algorithm**

Percent of replicates where one GP algorithm (y-axis) outperformed another GP algorithm (x-axis) for predicting each species/trait combination.

**Table S1. Hyperparameters examined in grid search**

**Table S2. Results of ANOVA on impact of each hyperparameter for each GP algorithm on model performance for height in each species**

**Table S3. Standard deviation of model performance across replicates for ANN and seeded ANN models for predicting height in each species.**

**Table S4. Full benchmark predictive performance results using all GP algorithms on all species/trait combinations**

**Table S5. Summary of GP algorithm performance across benchmark analysis Table S6. Publications included in analysis of GP algorithm comparisons**

