## Supplementary figures and images for "Benchmarking algorithms for genomic prediction of complex traits"

### Supplemental Figures

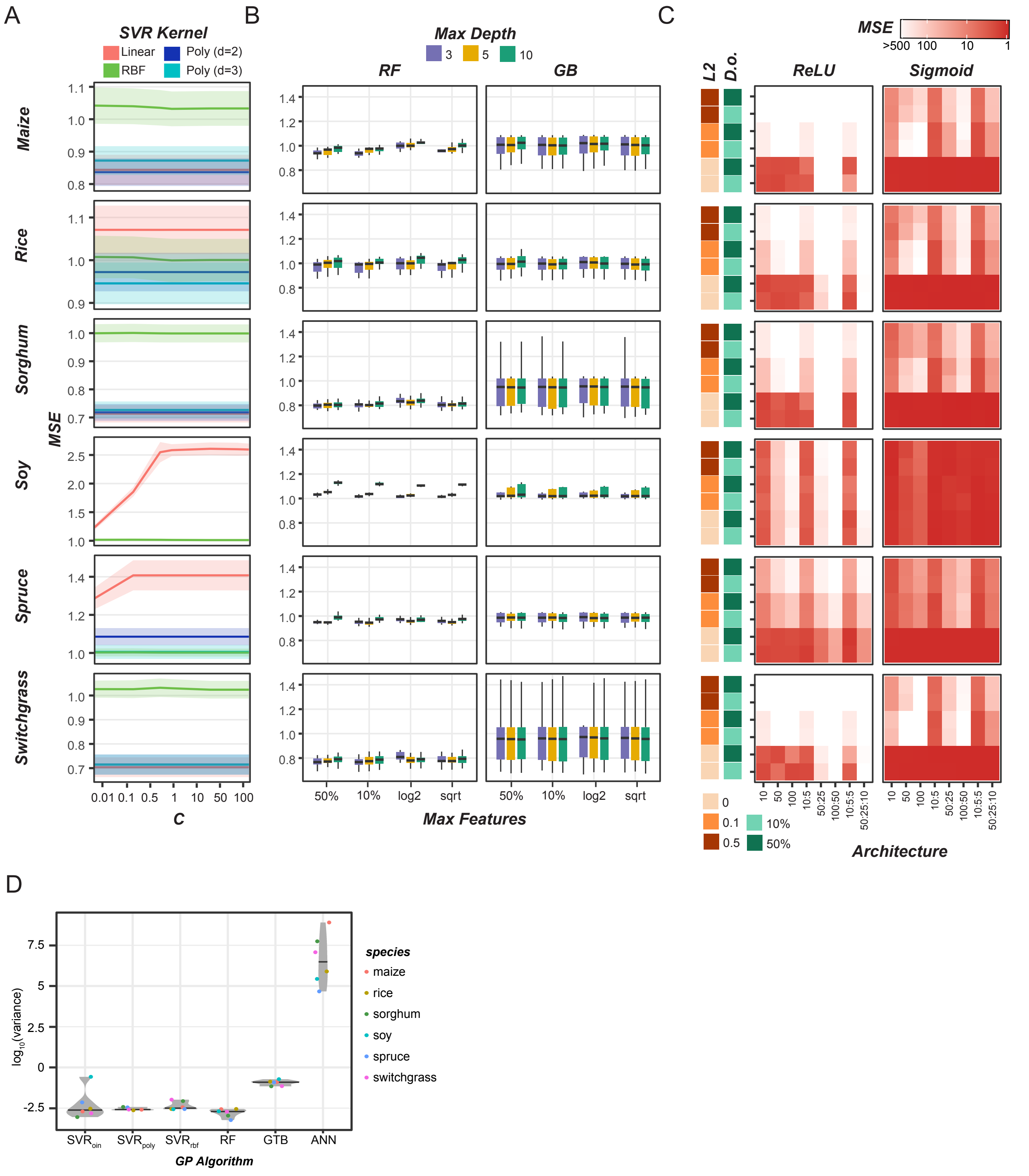

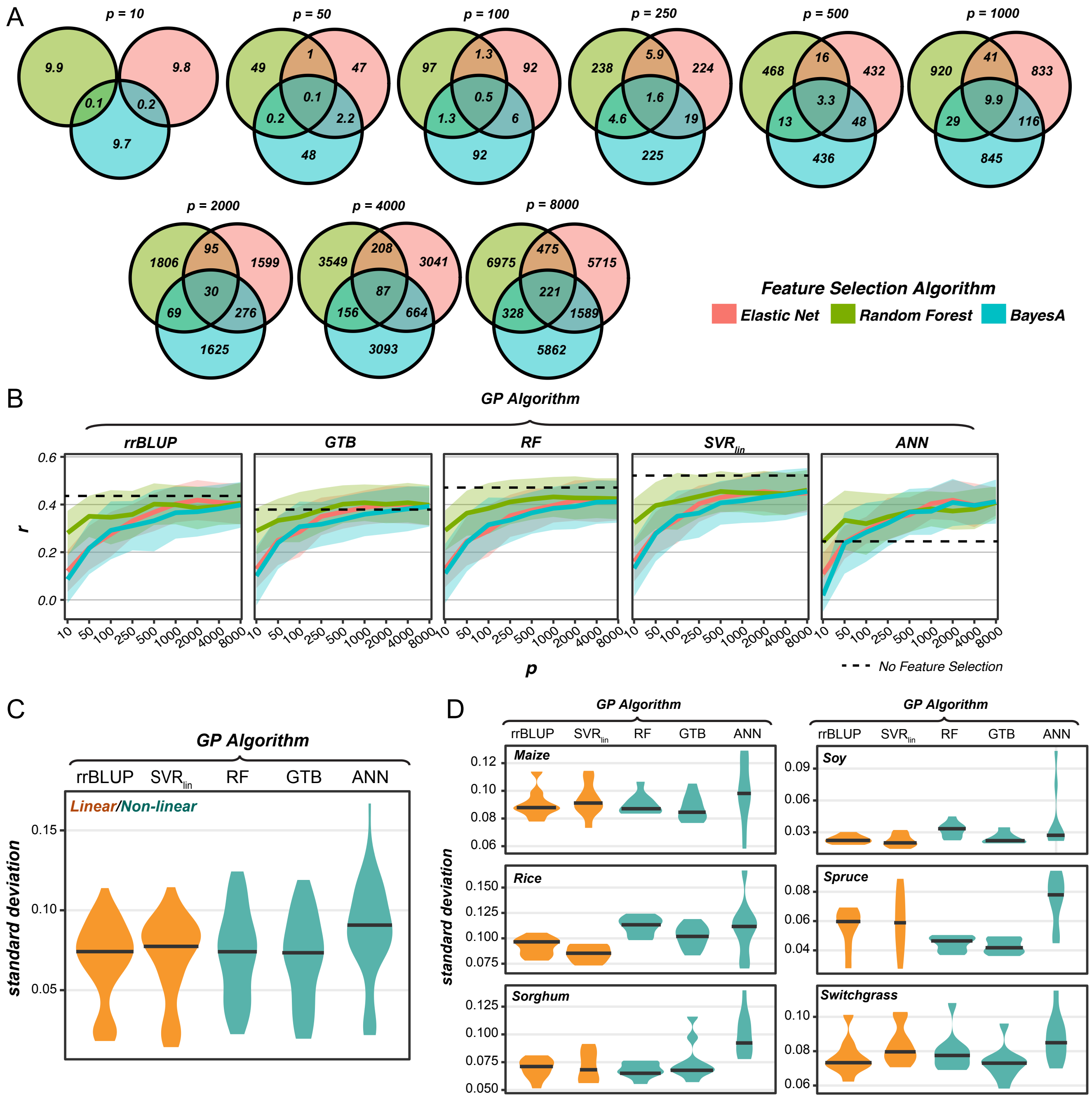

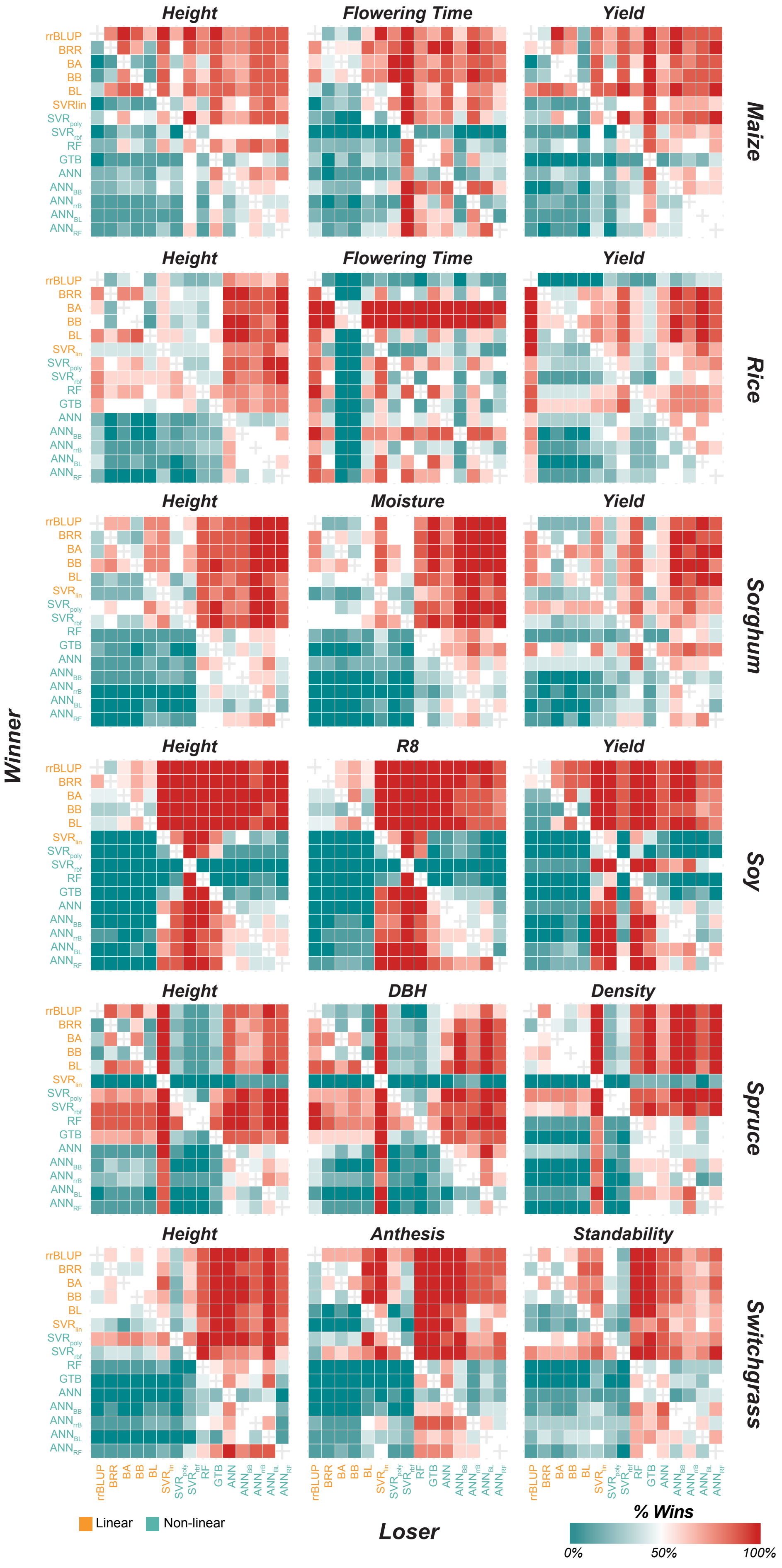
